# Coralline companions: Exploring microbes in coral ecosystems

**DOI:** 10.1101/2024.08.08.607119

**Authors:** Daizy Bharti, Arnab Ghosh, Santosh Kumar

## Abstract

Coral reefs exhibit remarkable resilience to natural disturbances but face challenges due to global changes and emerging diseases. In the present study, we recorded a total of 51 ciliates and over 400 bacteria associated with healthy and diseased corals from the Ramsar site Gulf of Mannar Marine National Park, India, based on morphology, silver staining, and Next Generation Sequencing. Our study contributes to the growing understanding of ciliate diversity and provides insights into the dynamics of ciliate-associated coral diseases, towards developing effective biocontrol agents for coral bleaching and conservation strategies to safeguard coral reef ecosystems.

---

In the absence of human influence, coral reefs demonstrate a notable capacity for rapid reassembly following natural disturbances (Connell et al., 1997). However, there is evidence of a gradual decline in the regeneration ability of coral reefs due to global changes, such as ocean warming, marine heat waves (MHWs), sea level rise, altered storm patterns, precipitation variations, shifts in ocean currents, ocean acidification, overexploitation of reef resources, nutrient enrichment from pollution, and coastal development (Coles and Brown, 2003; Bourne et al., 2009; Ainsworth et al., 2010). Additionally, the emergence of various diseases has become a significant contributing factor to coral destruction. These diseases negatively impact coral reproduction and growth rates, thereby altering community structure, species diversity, and the abundance of organisms associated with reefs (Loya et al., 2001). The primary microbial assemblages involved include bacteria, fungi, viruses, and protozoa, particularly ciliates (Sokolow, 2009; Sweet and Bythell, 2012). Nutrients provided by these associated microbes are utilized by corals for reef formation through the deposition of calcium carbonate and the secretion of carbon-rich mucus (Webster and Reusch, 2017).

In coral reefs, ciliates, particularly mixotrophs and heterotrophs, are attracted by the mucus secreted by corals according to their food preferences, thus playing a crucial role in nutrient cycling. This attraction enhances nitrogen, phosphorus, and extractable sulfur levels, ultimately promoting the growth of beneficial microbes associated with corals (Ravindran et al., 2003). However, certain microscopic eukaryotes are implicated in several coral diseases characterized by macroscopic lesions, such as White Syndrome (WS), Brown Band Disease (BrB), Porites White Patch Syndrome (PWPS), White Plaque (WP), White Band (WB), Skeletal Eroding Band (SEB), Caribbean Ciliate Infection (CCI), Caribbean Yellow Band Disease, Black Band Disease, among others (Sweet and Séré, 2016; Ravindran et al., 2023), thereby increasing the risk of coral mortality (Sweet and Bythell, 2012).

SEB, caused by *Halofolliculina corallasia*, marked the first ciliate-associated coral disease (Winkler et al., 2004). Subsequently, *Porpostoma guamense* was linked to BrB, affecting over 26 Caribbean coral species and termed CCI (Lobban et al., 2011). WS and other diseases like WBD show diverse ciliate communities, suggesting opportunistic feeding on decaying tissue (Sweet and Bythell, 2012; Randall et al., 2014; Sweet et al., 2014). Recent studies support this, especially for WBD and WS (Sweet et al., 2014; Sweet and Bythell, 2015).

Although the precise mechanism by which ciliates infect corals remains poorly understood, a crucial initial step in comprehending their role in coral diseases involves assessing the ciliate community associated with various disease states on a global scale. In India, four species (*Euplotes* sp., *Cohnilembus* sp., *Uronema marinum*, and *Holosticha* sp.) have been identified thus far that may potentially be associated with diseases in coral species from the Gulf of Mannar Marine National Park (GoM) (Ravindran et al., 2023).

Ciliates typically play a significant role in maintaining microbial biomass (Vargas and Hattori, 1990) and bacterial community structure, thereby exerting control over both benthic and pelagic food webs across various microhabitats. To delve deeper into this interaction, we examined the ciliate and bacterial communities associated with both healthy and diseased corals from the GoM. Our investigation identified a total of 51 ciliate species and over 400 bacterial species, offering valuable baseline insights, particularly regarding ciliates. Sampling efforts covered a large area (>150 km) of the GoM, spanning more than 15 small islands. Among the 51 recorded ciliates, 22 were previously known to be associated with coral diseases (Fig. 1,2; Supplemental Fig. S1-3, Table 1, 2). Surprisingly, 80% (i.e., 21 species, See Supplemental Table 2) of the 27 ciliate species known to be linked to coral diseases worldwide were found in the GoM. Additionally, we identified 40 ciliate species associated with diseased corals, 22 associated with healthy corals, and 11 associated with both diseased and healthy corals. This data underscores the complexity of interactions between ciliates and corals, as the ciliate community comprises various trophic groups, including bacterivores, herbivores, carnivores, histophages, commensals, parasites, and mixotrophs. Mixotrophic ciliates have been observed to proliferate significantly during marine heatwave events. It’s known that marine ciliates inhabiting vast, thermally stable oceans are highly adapted to ambient temperatures, with lower environment-specific activation energies compared to those in more variable freshwater environments (Lukić et al., 2022). This suggests that marine heatwaves could potentially disrupt ciliate compositions and consequently impact coral health.

**Fig. 1.**
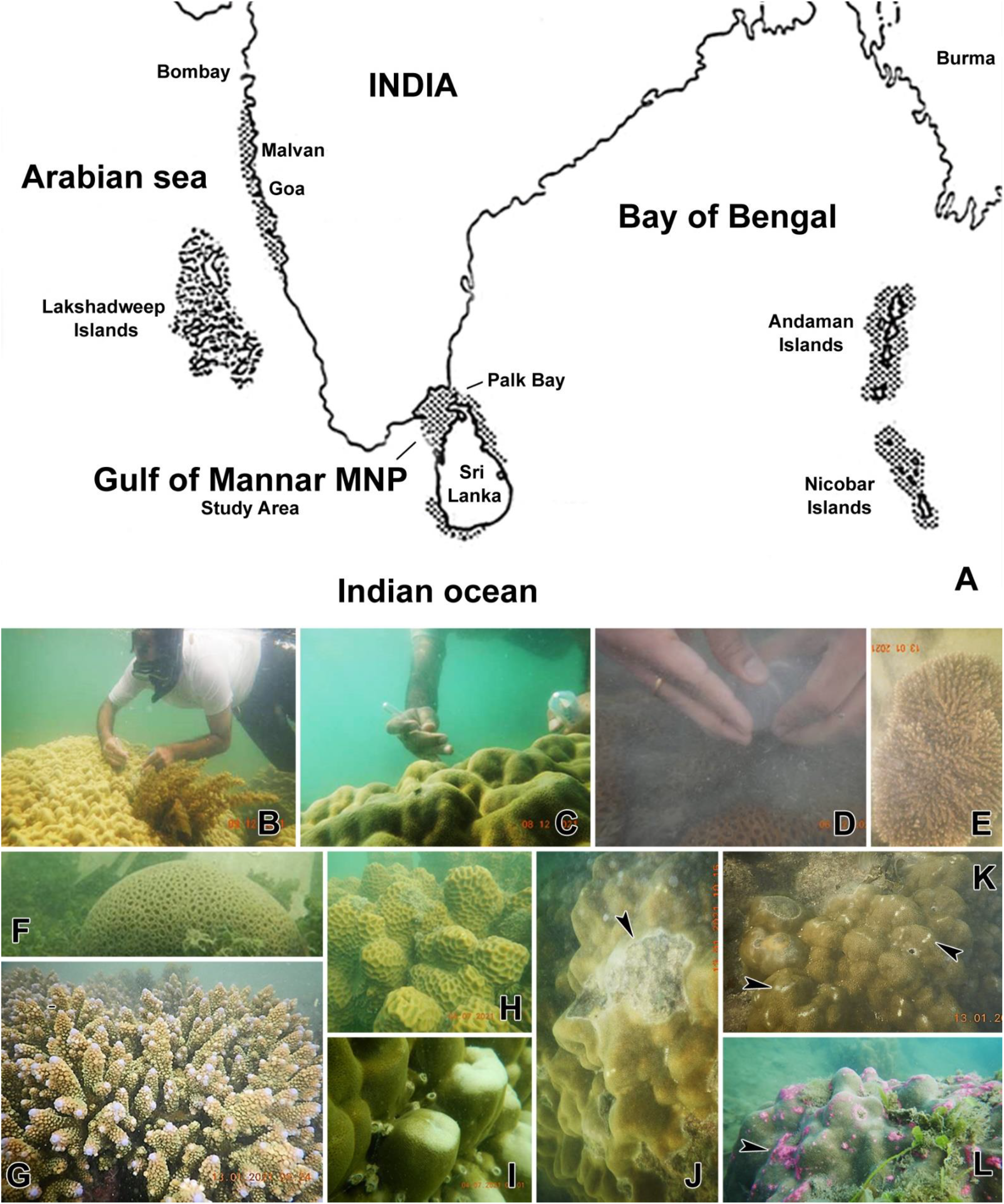
**(A)**Map showing the distribution of corals in India and the study site, i.e., Gulf of Mannar Marine National Park, India (map modified from D.O.D and S.A.C. 1997). **(B-D)** The author, Santosh Kumar, (B) during sample collection. Samplings were performed by collecting water and sediments with the help of pipettes by gently brushing the healthy corals surfaces and by carefully scratching the disease sites in corals. **(E-I)** Live corals. **(J-L)** Diseased corals (arrowheads point to the lesions).

**Fig. 2.**
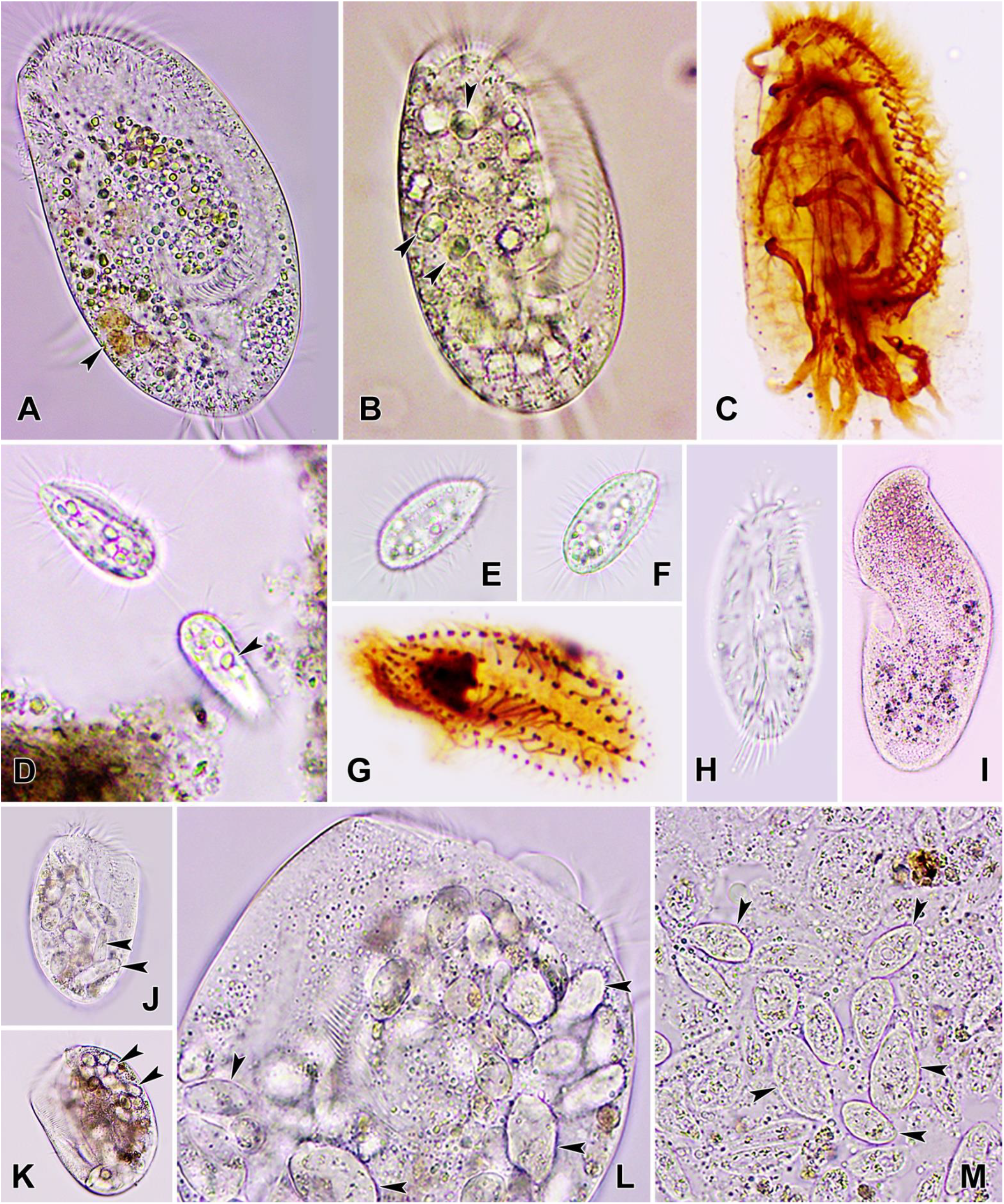
Photomicrograph of ciliates species associated with corals of Gulf of Mannar Marine National Park, India. **(A-C)** Specimens of *Moneuplotes vannus* from live (A, B) and after silver impregnation (C). **(D-G)** Specimens of *Uronema heteromarinum* from live (D-F) and after Silver impregnation (G). Specimens in D showing feeding around dead tissue and sediments. **(H)** *Holosticha diademata*, reported to be feeding on tissue of corals and zooxanthellae alga. **(I)** *Metopus contortus*, representative of anaerobic conditions. **(J-M)** Specimens of *Euplotes* sp. from live (large morphs). A possible bio-control agent for controlling *Uronema heteromarinum* infections in corals. Arrowheads points to the Zooxanthellae algae as food vacuole in the cytoplasm.

We noted the frequent presence and wide distribution of *Moneuplotes vannus* and *Uronema heteromarinum* across our samples, spanning nearly 150 km, indicating their tolerance and adaptability to adverse conditions (Fig. 2). Furthermore, we observed a *Euplotes* species voraciously feeding on specimens of *U. heteromarinum*, which is implicated in diseases such as White Syndrome (WS), Porites White Patch Syndrome (PWPS), and White Plaque (WP), suggesting its potential as a biocontrol agent for the latter (Fig. 2). Interestingly, this *Euplotes* species exhibited two types of morphs: small and large. The small morphs formed when the cells of *Euplotes* fed on bacteria and algae, whereas the large morphs (Fig. 2J-M) resulted from voracious feeding on *U. heteromarinum*. This indicates the wide food preference of this species and their ability to switch to *U. heteromarinum* as a food source. This adaptability shows significant potential for controlling *U. heteromarinum* populations, which have a large distribution in the Gulf of Mannar and are known to be associated with disease progression (bleaching) in corals (Fig. 2).

The presence of *Metopus* species indicates anoxic environments, due to algae sedimentation observed during the sampling events, suggesting that anaerobic ciliates also play specific roles in coral health. Additionally, the identification of *Holosticha diademata* suggests the presence of disease progression by ciliates, as this species has been reported to invade coral tissue and feed on Symbiodiniaceae, leading to increased disease lesions. However, the roles of these microbes need to be further investigated using Koch’s postulates to confirm whether they cause disease or are merely associated with lesion progression.

Analysis of bacterial communities based on Next Generation Sequencing (NGS) from healthy and diseased corals revealed the presence of over 400 different bacterial species (Fig. S3). Notably, 10% of the bacterial species present in diseased corals belonged to the family Vibrionaceae, known to cause diseases in corals and tested using Koch’s postulates (Séré et al., 2015). The presence of 40 ciliate species suggests a potential interaction between these groups, although it remains unclear whether ciliates are controlling bacterial populations or exacerbating disease progression.

One major challenge in ciliate research globally is their minute size, complex methodologies, and a scarcity of experts in this field (Foissner et al., 2005). In this context, the identification of 51 species, 40 of which are associated with coral disease, contributes to the expanding catalogue of ciliates reported worldwide, with about 28 species known to cause infections in corals (Supplemental Fig. S1-3, Table 1, 2). Furthermore, the observation of Symbiodiniaceae alga within the cytoplasm of many ciliates recorded in our study indicate that ciliates may opportunistically feed on corals already infected by other pathogens, consuming damaged tissue containing Symbiodiniaceae, thereby disrupting coral reassembly properties and potentially threatening surrounding biodiversity.

It is hypothesized that coral species could serve as umbrella species, providing protection for all other species directly or indirectly associated with coral species or structures. Nevertheless, further marine ecosystem surveys may strengthen this assertion and increase our understanding of ciliate diversity and dynamics of interaction. In conclusion, the present work supplements the efforts to enhance our knowledge of ciliate diversity and understand complex interactions affecting coral health. The study lays the groundwork for future research into the role of the ciliate community and their functional significance in relation to corals, as well as the search for ciliates as biocontrol agents for coral bleaching. Such efforts are crucial for coral conservation, especially at these times of MHWs occurrences worldwide, towards fostering the growth and development of healthy corals, and preserving the biological diversity dependent on them.

## Supplementary Material

Supplementary information is available at xx. The metagenomics data is being submitted to international repositories.

## Acknowledgments

The authors would like to express their gratitude to Dr. Dhriti Banerjee, Director of the Zoological Survey of India (ZSI), Kolkata, for support and encouragement. CSIR-India fellowship to Mr. Arnab Ghosh is duly acknowledged. Special thanks are extended to Dr. N. Marrimuthu, Scientist-E, ZSI, Kolkata, for his invaluable assistance and support in sample collection. The authors are also grateful to the forest department, Tamil Nadu, India for granting permission and offering support during the sampling process. This study is part of an internal program of the Zoological Survey of India, Kolkata, India, titled “*Studies on the ciliates and bacterial communities associated with corals based on morphology and molecular tools*” led by S. Kumar at ZSI, Kolkata.

## Author Contributions

Daizy Bharti: Conceptualization; Data curation; Formal analysis; Methodology; Writing -original draft; Writing - review & editing. Arnab Ghosh: Formal analysis; Methodology; Writing - original draft; Writing - review & editing. Santosh Kumar: Conceptualization; Investigation; Methodology; Project administration; Supervision; Validation; Visualization; Writing - original draft; Writing - review & editing.

