## Supplementary file for "Coralline companions: Exploring microbes in coral ecosystems"

### Supplementary materials

#### Materials & Methods

Sampling was conducted in the Gulf of Mannar Marine National Park, India, spanning a stretch from Tuticorin to Rameswaram, covering approximately 150 km in Tamil Nadu, India, and including nearly 15 Islands. Overall 137 samples were collected on three surveys conducted, i.e., 03.01.2021 to 17.01.2021 with collection of 52 samples; 04.12.2021 to 10.12.2021 with collected 48 samples; and 04.07.2022 to 10.07.2022 with collection of 37 samples. Samples were collected through skin diving. Collection involved gently extracting mucus from live corals and lesions from deceased corals using pipettes and syringes, which were then deposited into tubes. Subsequently, the 137 samples collected during the studies were transported to the laboratory for processing. To document species characteristics and for identification, live observations were conducted, and silver staining was performed to study morphology (Bharti and Kumar, 2023). Live observations were facilitated using an Olympus (CX 43) microscope with bright-field illumination. The protargol staining method, as described by Kamra and Sapra (1990) and Foissner (1991), with certain modifications, was employed to reveal the infraciliature. For analysis of the bacterial community, samples were obtained from both healthy and diseased corals. Sequencing were performed at LCGC sequencing services, Hyderabad, India. DNA extraction for metagenomic analyses was carried out using commercially available kits from QIAGEN, ZYMO RESEARCH, and ThermoFisher. PCR amplification of the V3-V4 region of the 16S gene was performed using TAQ Master MIX, comprising High-Fidelity DNA Polymerase, 0.5mM dNTPs, 2mM $MgCl_2$, and PCR enzyme buffer. The primers used were V13F (5′ AGAGTTTGATGMTGGCTCAG 3'). Raw data quality control was conducted using FASTQC and MULTIQC, followed by trimming of adapters and low-quality reads using TRIMGALORE. The trimmed reads were subsequently processed, including merging of paired-end reads, removal of chimeras, and abundance calculation and estimation correction utilizing QIIME, MOTHUR, KRAKEN, and BRACKEN workflows.

**Supplemental Table 1.** Coral associated ciliates identified from the Gulf of Mannar Marine National Park, India.

| S. No. | Present study | Associated Corals species | Sampled from diseased corals | Sampled from healthy corals | Observed in samples |
|---|---|---|---|---|---|
| 1. | *Aegeria* sp. | *Dipsastraea* sp. | - | + | 01 |
| 2. | *Aspidisca steinii* | *Acropora* sp., *Goniastrea* sp., *Porites* sp. | + | - | 05 |
| 3. | *Chaenea vorax* | *Porites* sp. | + | - | 01 |
| 4. | *Cohnilembus verminus* | *Porites* sp. | + | - | 05 |
| 5. | *Condylostoma magnum* | *Favia* sp. | + | - | 01 |
| 6. | *Dysteria derouxi* | *Porites* sp. | + | - | 01 |
| 7. | *Euplotes minuta* | *Favia* sp., *Goniastrea* sp., *Porites* sp. | + (*Favia*, *Porites*) | + (*Goniastrea*) | 11 |
| 8. | *Euplotes* sp. | *Favia* sp. | + | + | 01 |
| 9. | *Frontonia canadensis* | *Favia* sp. | + | | 01 |
| 10. | *Glauconema trihymene* | *Dipsastraea* sp., *Porites* sp. | - | + | 02 |

| | | | | | |
|---|---|---|---|---|---|
| 11. | *Hartmannula derouxi* | *Leptoria* sp., *Porites* sp. | + (*Porites*) | + (*Leptoria*) | 04 |
| 12. | *Hemigastrostyla enigmatica* | *Favia* sp., *Porites* sp. | + (*Porites*) | + (*Favia*) | 02 |
| 13. | *Holosticha diademata* | *Acropora* sp., *Goniastrea* sp. | + | - | 02 |
| 14. | *Lacrymaria coronata* | *Favia* sp. | - | + | 01 |
| 15. | *Litonotus pictus* | *Porites* sp. | + | - | 01 |
| 16. | *Mesodinium* sp. | *Favia* sp. | | + | 01 |
| 17. | *Metacystis stariata* | *Acropora muricata*, *Porites* sp. | + | - | 02 |
| 18. | *Metanophrys* sp. | *Favia* sp. | - | + | 02 |
| 19. | *Metopus contortus* | *Porites* sp. | + | - | 02 |
| 20. | *Metopus* sp. | *Favia* sp. | + | - | 01 |
| 21. | *Moneuplotes vannus* | *Acropora* sp., *Astreopora* sp., *Favites* sp., *Goniastrea* sp., *Porites* sp., *Platygyra* sp. | + (*Acropora*, *Porites*) | + (*Acropora*, *Astreopora*, *Favites*, *Goniastrea*, *Porites*, *Platygyra*) | 16 |
| 22. | *Nothoholosticha flava* | *Favia* sp., *Porites* sp. | + | - | 03 |
| 23. | *Oxytricha lithofera* | *Acropora* sp. | + | - | 01 |
| 24. | *Oxytricha sp.* | *Goniastrea* sp. | | + | 01 |
| 25. | *Paramesanophrys* sp. | *Porites* sp. | + | - | 01 |
| 26. | *Paramesanophrys typicus* | *Montipora* sp., *Porites* sp. | + (*Porites*) | + (*Montipora*) | 02 |
| 27. | *Paranophrys* sp. 1 | *Favites* sp. | - | + | 02 |
| 28. | *Paranophrys* sp. 2 | *Favites* sp. | - | + | 01 |
| 29. | *Parauronema longum* | *Acropora* sp., *Porites* sp. | + | - | 02 |
| 30. | *Pelagostrobilidium minutum* | *Favia* sp. | + | - | 01 |
| 31. | *Peritromus faurei* | *Porites* sp. | + | - | 01 |
| 32. | *Porpostoma guamense* | *Goniastrea* sp. | | + | 01 |
| 33. | *Philaster lucinda* | *Acropora* sp., *Porites* sp. | + | - | 02 |
| 34. | *Pleuronema orientale* | *Acropora* sp. | + | - | 02 |
| 35. | *Pleuronema wiackowskii* | *Porites* sp. | + | - | 01 |
| 36. | *Protocruzia adherens* | *Goniastrea* sp., *Porites* sp. | + (*Goniastrea*, *Porites*) | +(*Goniastrea*, *Porites*) | 10 |
| 37. | *Protocruzia contrax* | *Acropora* sp., *Goniastrea* sp., *Porites* sp. | + (*Acropora*, *Porites*) | + (*Goniastrea*) | 03 |
| 38. | *Protocruzia* sp. | *Porites* sp. | + | - | 01 |
| 39. | *Pseudocohnilembus hargisi* | *Porites* sp. | + | - | 01 |
| 40. | *Pseudokeronopsis flava* | *Favia* sp. | + | - | 01 |
| 41. | *Strombidium caudispina* | *Acropora* sp., *Favia* sp., *Goniastrea* sp. | + (*Acropora*, *Goniastrea*) | + (*Favia*) | 03 |
| 42. | *Strombidium* sp. | *Goniastrea* sp., *Porites* sp. | + | - | 02 |
| 43. | *Strombidium sulcatum* | *Goniastrea* sp. | + | - | 02 |
| 44. | *Trachelocerca* sp. | *Porites* sp. | + | + | 02 |
| 45. | *Trachelostyla pediculiformis* | *Goniastrea* sp. | + | - | 01 |
| 46. | *Trochiliopsis* sp. | *Acropora* sp., | - | + | 01 |
| 47. | *Uronema elegans* | *Favia* sp., *Porites* sp. | + | - | 02 |
| 48. | *Uronema heteromarinum* | *Acropora* sp., *Dipsastraea* sp., *Favites* sp., | + (*Acropora*, *Goniastrea*, | + (*Acropora*, *Dipsastraea*, | 18 |

| | | | | | |
|---|---|---|---|---|---|
| | | *Goniastrea* sp., *Leptoria* sp., *Porites* sp., *Pocillopora* sp. | *Leptoria*, *Porites*, *Pocillopora*) | *Favites*, *Goniastrea*, *Leptoria*, *Porites*) | |
| 49. | *Uronychia setigera* | *Favia* sp., *Porites* sp. | + | - | 04 |
| 50. | *Zosterodasys* sp. | *Favia* sp. | - | + | 01 |
| 51. | *Pseudovorticella* sp. | *Acropora* sp. | + | - | 01 |

**Supplemental Table 2.** Ciliate species identified in the present study from the Gulf of Mannar marine National Park, India, that are known to be associated with diseases in corals (refer Sweet and Séré 2015; Ravindran et al. 2023).

| S. No. | Species | Reported from coral diseases |
|---|---|---|
| *1.* | *Aspidisca steinii* | White Syndrome (WS) |
| *2.* | *Chaenea vorax* | White Syndrome (WS), White Plaque (WP), White Band (WB), Skeletal Eroding Band (SEB), Black Band Disease (BBD), |
| *3.* | *Cohnilembus verminus* | Feeding on tissue lesions |
| *4.* | *Condylostoma magnum* | Skeletal Eroding Band (SEB) |
| *5.* | *Dysteria derouxi* | White Syndrome (WS), Porites White Patch Syndrome (PWPS), White Plaque (WP), White Band (WB), Skeletal Eroding Band (SEB) |
| *6.* | *Euplotes* sp. | White Syndrome (WS), Brown Band Disease (BRB) |
| *7.* | *Frontonia canadensis* | Tissue lesions |
| *8.* | *Glauconema trihymene* | White Syndrome (WS), Brown Band Disease (BRB), White Band (WB), |
| *9.* | *Hartmannula derouxi* | White Syndrome (WS) |
| *10.* | *Hemigastrostyla enigmatica* | White Syndrome (WS) |
| *11.* | *Holosticha diademata* | White Syndrome (WS), Brown Band Disease (BRB), Porites White Patch Syndrome (PWPS), White Plaque (WP), Skeletal Eroding Band (SEB), Black Band Disease (BBD), Caribbean Yellow Band Disease (CYBD) |
| *12.* | *Litonotus pictus* | White Syndrome (WS) |
| *13.* | *Moneuplotes vannus* | White Syndrome (WS), Brown Band Disease (BRB) |
| *14.* | *Nothoholosticha flava* | White Syndrome (WS), White Plaque (WP) |
| *15.* | *Philaster lucinda* | Brown Band Disease (BRB), Porites White Patch Syndrome (PWPS), White Plaque (WP), White Band (WB), Skeletal Eroding Band (SEB), Caribbean Ciliate Infections (CCI), Black Band Disease, |
| *16.* | *Porpostoma guamense* | Brown Band Disease (BRB), White Syndrome (WS) |
| *17.* | *Protocruzia adherens* | White Syndrome (WS), White Plaque (WP), White Band (WB), Black Band Disease (BBD), Caribbean Yellow Band Disease (CYBD) |
| *18.* | *Pseudokeronopsis flava* | White Syndrome (WS), Brown Band Disease (BRB), White Band (WB), |
| *19.* | *Strombidium* sp. | White Syndrome (WS), Brown Band Disease (BRB), Porites White Patch Syndrome (PWPS), White Plaque (WP), White Band (WB), Skeletal Eroding Band (SEB) |
| *20.* | *Uronema heteromarinum* | White Syndrome (WS), Porites White Patch Syndrome (PWPS), White Plaque (WP) |
| *21.* | *Uronychia setigera* | White Syndrome (WS), Brown Band Disease (BRB) |

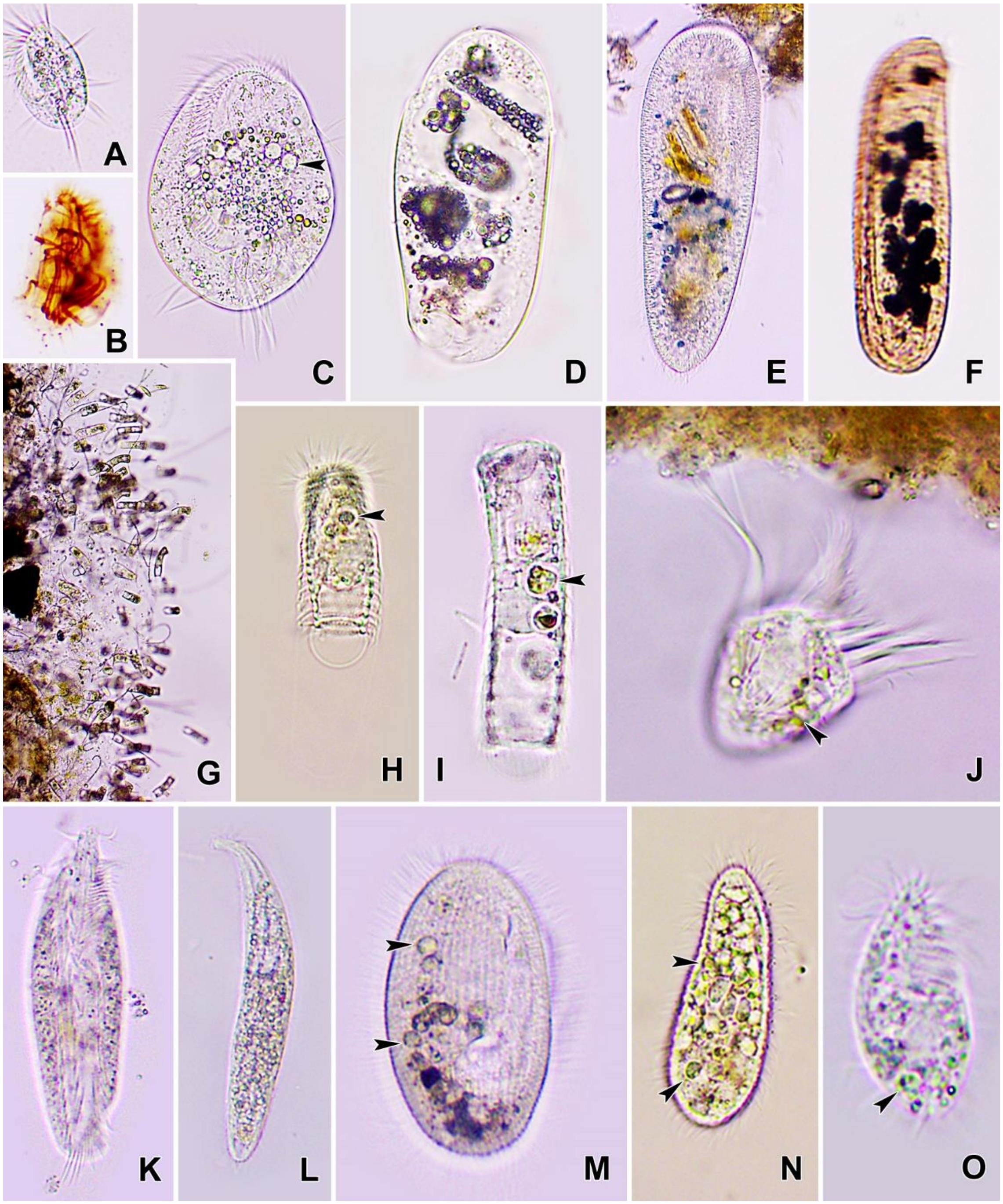


**Figure S1.** Ciliates isolated from corals of the Gulf of Mannar Marine National Park, India. **(A-C)** *Euplotes minuta* from live (A,C) and after protargol impregnation (B). **(D)** *Zosterodasys* sp. **(E)** *Frontonia canadensis*. **(F)** *Dysteria derouxi*. **(G-I)** *Metacystis stariata*. **(J)** *Strombidium* sp. **(K)** *Hemigastrostyla enigmatica*. **(L)** *Trachelocerca* sp. **(M)** *Pleuronema orientale.* **(N)** *Hartmannula derouxi*. **(O)** *Protocruzia adherens*. Arrowheads points to the Zooxanthellae algae as food vacuole in the cytoplasm.

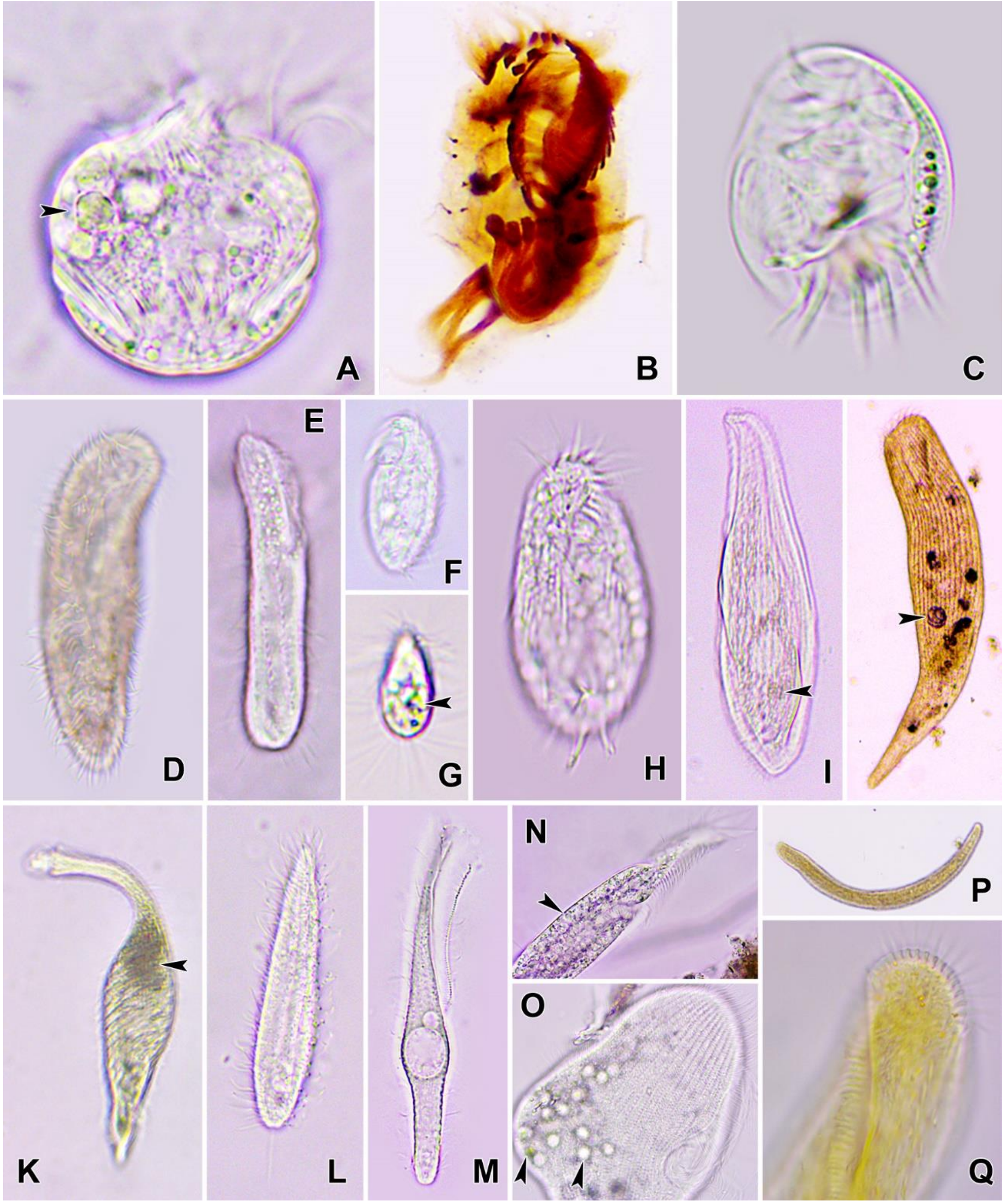


**Figure S2.** Ciliates isolated from corals of the Gulf of Mannar Marine National Park, India. **(A)** *Strombidium sulcatum.* **(B)** *Uronychia setigera*. **(C)** *Aspidisca steinii*. **(D)** *Pseudokeronopsis flava*. **(E)** *Pseudocohnilembus hargisi*. **(F)** *Protocruzia contrax*. **(G)** *Mesodinium* sp. **(H)** *Oxytricha lithofera*. **(I)** *Litonotus pictus*. **(J)** *Condylostoma magnum*. **(K)** *Lacrymaria coronata*. **(L)** *Chaenea vorax*. **(M)** *Cohnilembus verminus*. **(N)** *Trachelostyla pediculiformis*. **(O)** *Peritromus faurei*. **(P,Q)** *Nothoholosticha flava.* Arrowheads points to the Zooxanthellae algae as food vacuole in the cytoplasm.

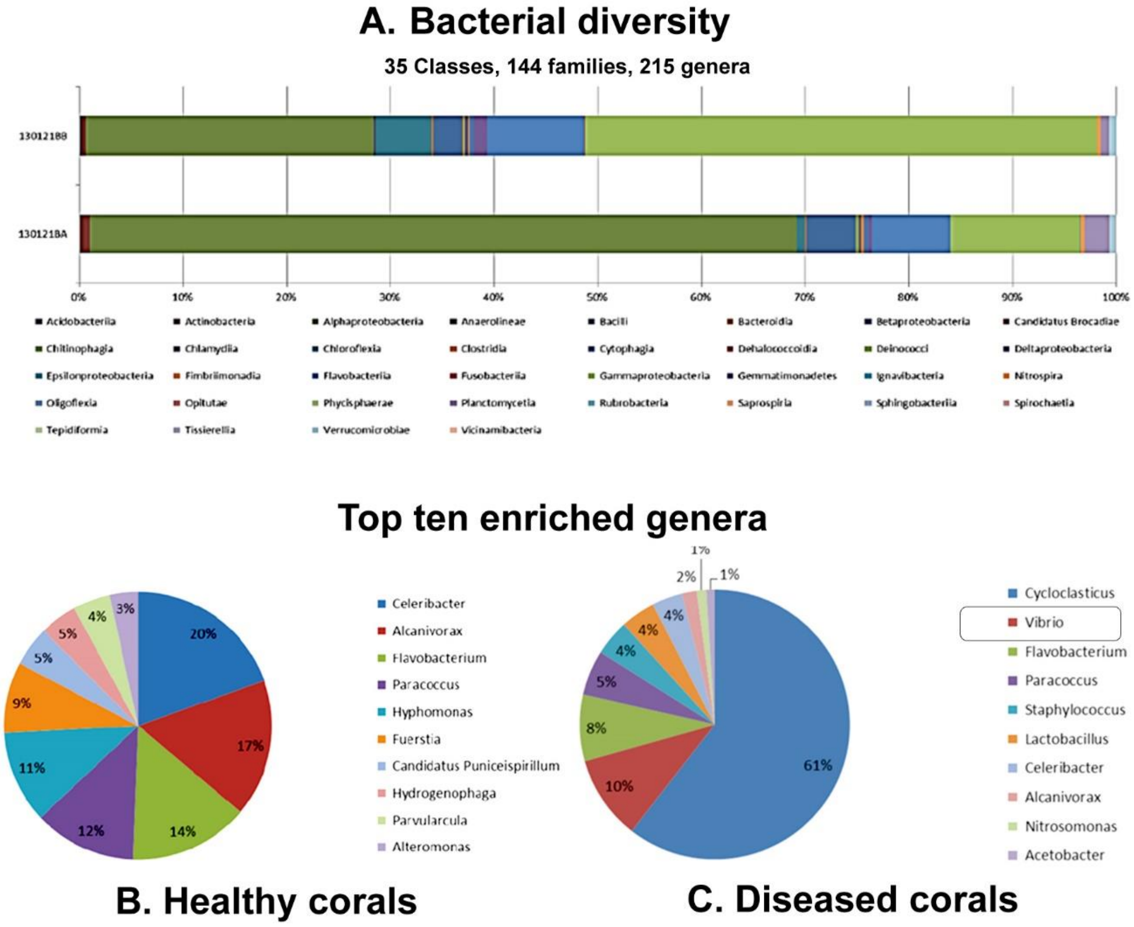


**Figure S3. (A-C)** Showing bacterial diversity in healthy (sample code: 130121BA) and diseased corals (sample code: 120121BB). Note the presence of vibrio (10%) in diseased coral sample based on data from top ten enriched genera.

#### Supplementary materials references


Bharti, D., Kumar, S., 2023. Description of a new oxytrichid ciliate, *Oxytricha buxai* n. sp. and redescription of *O. quadricirrata* Blatterer and Foissner, 1988 based on morphology and 18S rDNA analyses. Eur. J. Protistol. 88, 125959.

DOD, SAC., 1997. Coral reef maps of India, Department of Ocean Development and Space Application Centre, Ahmedabad, India.

Foissner, W., 1991. Basic light and scanning electron microscopic methods for taxonomic studies of ciliated protozoa. Eur. J. Protistol. 27, 313–330.

Kamra, K., Sapra, G.R., 1990. Partial retention of parental ciliature during morphogenesis of the ciliate Coniculostomum monilata (Dragesco and Njiné, 1971) Njiné, 1978 (Oxytrichidae, Hypotrichida). Eur. J. Protistol. 25, 264–278.
